# Deep Docking - a Deep Learning Approach for Virtual Screening of Big Chemical Datasets

**DOI:** 10.1101/2019.12.15.877316

**Authors:** Francesco Gentile, Vibudh Agrawal, Michael Hsing, Fuqiang Ban, Ulf Norinder, Martin E. Gleave, Artem Cherkasov

## Abstract

Drug discovery is an extensive and rigorous process that requires up to 2 billion dollars of investments and more than ten years of research and development to bring a molecule “from bench to a bedside”. While virtual screening can significantly enhance drug discovery workflow, it ultimately lags the current rate of expansion of chemical databases that already incorporate billions of purchasable compounds. This surge of available small molecules presents great opportunities for drug discovery but also demands for faster virtual screening methods and protocols. In order to address this challenge, we herein introduce *Deep Docking* (*D*^*2*^) - a novel deep learning-based approach which is suited for docking billions of molecular structures. The developed *D*^*2*^-platform utilizes quantitative structure-activity relationship (QSAR) based deep models trained on docking scores of subsets of a large chemical library (Big Base) to approximate the docking outcome for yet unprocessed molecular entries and to remove unfavorable structures in an iterative manner. We applied *D*^*2*^ to virtually screen 1.36 billion molecules form the ZINC15 library against 12 prominent target proteins, and demonstrated up to 100-fold chemical data reduction and 6,000-fold enrichment for top hits, without notable loss of well-docked entities. The developed *D*^*2*^ protocol can readily be used in conjunction with any docking program and was made publicly available.

---

Drug discovery is an expensive and time-demanding process that faces many challenges, including low hit rates of high-throughput screening approaches among many others^1–2^. Methods of computer-aided drug discovery (CADD) can significantly speed up the pace of screening and help improving hit discovery rates^3^. Molecular docking is routinely used to process virtual libraries containing millions of molecular structures against a variety of drug targets with known three-dimensional structures. Thus, the ongoing explosive expansion of chemical databases represents great opportunities for virtual screening (VS) approaches in general and for docking in particular, but also poses entirely novel challenges. For instance, the widely-used ZINC library has grown from 700,000 entries in 2005^4^ to over 1.3 billion constituent molecular structures in 2019^5^, representing remarkable 1,000-fold increase. In a recent groundbreaking study by *Lyu et al*^6^, authors reported the largest docking experiment to date that involved 170 million molecular structures, which, however, still just nudges only one-tenth of the current ZINC. Thus, given the current state of docking protocols and computational resources available to CADD scientists, one can stipulate that modern docking campaigns can rarely exceed 0.1 billion molecules and that the current chemical space remains largely inaccessible to structure-based drug discovery.

One common approach to address this disparity is to filter large chemical collections to manageable drug-, lead-, fragment- and hit-like subsets (among others) using precomputed physicochemical parameters and drug-like criteria, such as molecular weight, volume, octanol-water partition coefficient, polar surface area, number or rotatable bonds, number of hydrogen bond donors and acceptors among many others^7^. While this approach can effectively reduce a Big Base to manageable subsets, many potentially useful compounds and novel or unconventional chemotypes (notably emerging from such large colllections^6^) could be lost. In order to take a full advantage of available and emerging ‘make-on-demand’ chemicals, it is essential to maximize the number of Big Base entries tangibly evaluated against a target of interest. It is also important to note, that conventional docking workflow is remarkably neglectful of negative results. Typical docking campaign relies on completing a full docking run and selecting an extremely narrow subset of favorably docked molecules (virtual hits) for future evaluation. Thus, the vast majority of docking data (both favorable and, especially unfavorable) is not being utilized in any way or form, while it could represent a very relevant, well-formatted and content-rich input for machine learning algorithms.

To confront this challenge, back in 2006 we introduced *Progressive Docking* - a hybrid docking/machine learning approach utilizing 3D ‘inductive’ QSAR (quantitative structure-activity relationship) descriptors^8–10^ to filter out molecules predicted to have unfavorable Glide docking scores^11^. While this method resulted in 3-4 fold enrichment of virtual hits (as several similar approaches reported more recently^12, 13^), the *Progressive Docking* did not gain a particular momentum. For that we could contemplate two possible reasons - on one hand, back in 2006 the ZINC database contained only ~1 million entries, which were amendable to conventional docking. On the other hand, back in those days only shallow learning approaches were available, which could not unlock the full power of synergy between docking and QSAR methodologies and could not take a full advantage of Big Data.

## RESULTS AND DISCUSSION

In the current study we have introduced several novel components into the original *Progressive Docking* workflow, including the use of fast-computed target-independent QSAR descriptors (such as 2D molecular fingerprint), the use of iterative and fast random sampling of the docking base (Big Base), and, principally, the use of Deep Learning (DL) to predict docking scores of yet unprocessed Big Base entries at each iteration step. As the result, the developed *Deep Docking D*^*2*^-pipeline achieved up to 100-fold reduction of Big Base, and up to 6,000-fold enrichment for the top-ranked hits, while avoiding significant loss of favorable virtual hits, as it will be discussed below.

### *D*^*2*^ pipeline

In essence, *D*^*2*^ pipeline (Figure 1) relies on the following consecutive steps:

a. For each entry of a Big Base (such as ZINC15), the standard set of ligand QSAR descriptors (such as molecular fingerprints) is computed;
b. A reasonably-sized training subset is randomly sampled from the Big Base and docked into the target of interest using conventional docking protocol(s);
c. The generated docking scores of the training compounds are then related to their 2D molecular descriptors through a DL model; a docking score cutoff (typically negative) is then used to divide training compounds in virtual hits (scoring below the cutoff) and non-hits (scoring above the negative cutoff);
d. The resulting QSAR deep model (trained on empirical docking scores) is then used to predict docking outcomes of yet unprocessed entries of the Big Base. A predefined number of predicted virtual hits are then randomly sampled and used for the training set augmentation;
e. Steps b) - d) are repeated iteratively until a predefined number of iterations is reached, or processed entries of a Big Base are converged.

**Figure 1.**
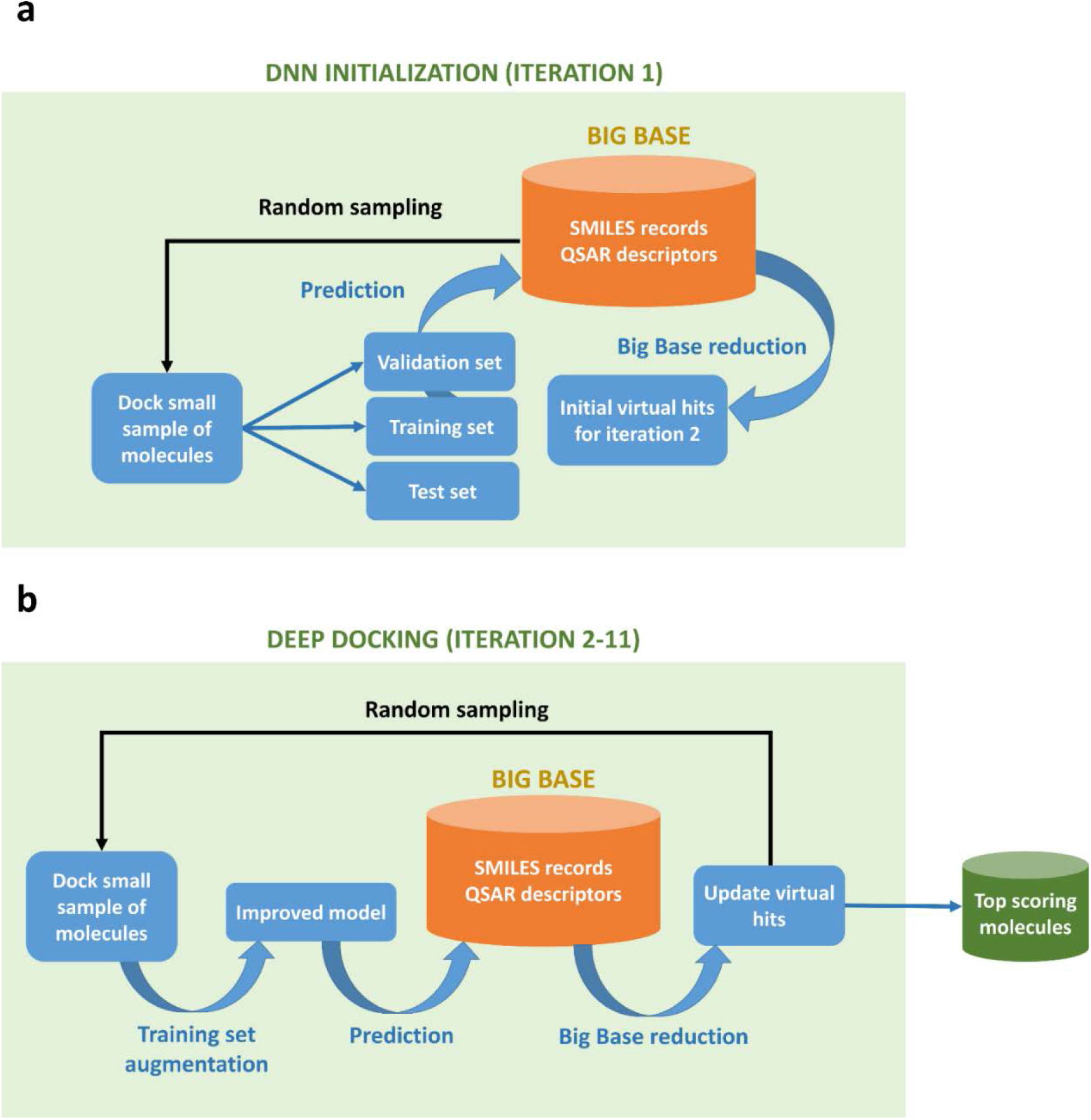
*Schematic of the* D^2^ *pipeline*. **a,** D^2^ initialization: a small sample of molecules is randomly extracted from a Big Base and docked to a target under consideration. The generated empirical docking scores are then used to train a Quantitative Structure-Activity Relationship (QSAR) deep model. The created QSAR solution is then used to predict docking outcome for the remainder of a Big Base, and to return predicted virtual hits required to start iteration 2. **b,** D^2^ screening: from iteration 2 onwards, the deep model gets gradually improved by augmenting the training set with randomly sampled QSAR-predicted virtual hits from the previous D^2^ iteration (which also get selected for actual docking). The cycle is repeated for a predefined number of iterations, after which D^2^ returns all top scoring molecules from a Big Base. This final library can be post-processed to remove residual low scoring entities. Alternatively, steps 2-11 can be carried out until the convergence of a Big Base.

The virtual hits recall from the Big Base (i.e. the percentage of actual virtual hits that is retrieved) is set implicitly in *D*^*2*^ through a probability threshold which is selected to include 90% of the actual virtual hits in the validation set. Then, the same threshold is applied to the independent test set and the recall of virtual hits is evaluated in order to assess model generalizability. If recalls of the validation and test sets are consistent with each other, the model is applied to all entries of the Big Base (more details can be found in the ‘Methods’ section). Although the recall values could be endorsed explicitly by using, for example, conformal predictors^13, 14^, we did not observe significant differences in the resulting performance of *D*^*2*^.

### Big Base sampling

The selection of a representative and balanced training set is a critical step of any modeling workflow. In the context on sampling a chemical space, a proper *D*^*2*^ training set should effectively reflect Big Base’s chemical diversity. It could be expected that enlarging the sampling size and pre-clustering the docking base would ultimately improve or even converge the chemical space coverage. On the other hand, it is currently not feasible to cluster billions of chemical structures in any way or form, and also the size of *D*^*2*^ training set (e.g. the amount of actual docking) would have a pivotal impact on a computational runtime and should be carefully controlled.

To establish an optimal sampling of ZINC15 base, we evaluated the relationship between the size of *D*^*2*^ training set and the corresponding means and standard deviations of the teste set recall values, reflecting the consistency of model’s performance and its generalizability. For that we evaluated 12 protein targets from four major drug-target families^15^, including nuclear receptors represented by Androgen Receptor (AR), Estrogen Receptor-alpha (ERα), and Peroxisome Proliferator-Activated Receptor γ (PPARγ). Kinases were represented by Calcium/Calmodulin-Dependent Protein Kinase Kinase 2 (CAMKK2), Cyclin-Dependent Kinase 6 (CDK6), Vascular Endothelial Growth Factor Receptor 2 (VEGFR2). G protein-coupled receptors included Adenosine A2A Receptor (ADORA2A), Thromboxane A2 Receptor (TBXA2R), Angiotensin II Receptor Type 1 (AT1R). Ion channels were represented by Nav1.7 sodium channel (Nav1.7), Gloeobacter Ligand-Gated Ion Channel (GLIC) and Gamma-aminobutyric acid receptor type A (GABAA) (more details about the selected targets are reported in Supplementary Table S1). For all 12 studied targets we investigated relationships between the sample size and resulting mean test set recall values, which appear to converge to 0.9 when the training set size approaches 1 million entries (Figure 2a). Similarly, it has been observed that the standard deviations also converge to 0 at about same ~1 million sample size (Figure 2b). Thus, we have set 1 million molecules as the standard sampling for *D*^*2*^ workflow (more details can be found in ‘Methods’ section).

**Figure 2.**
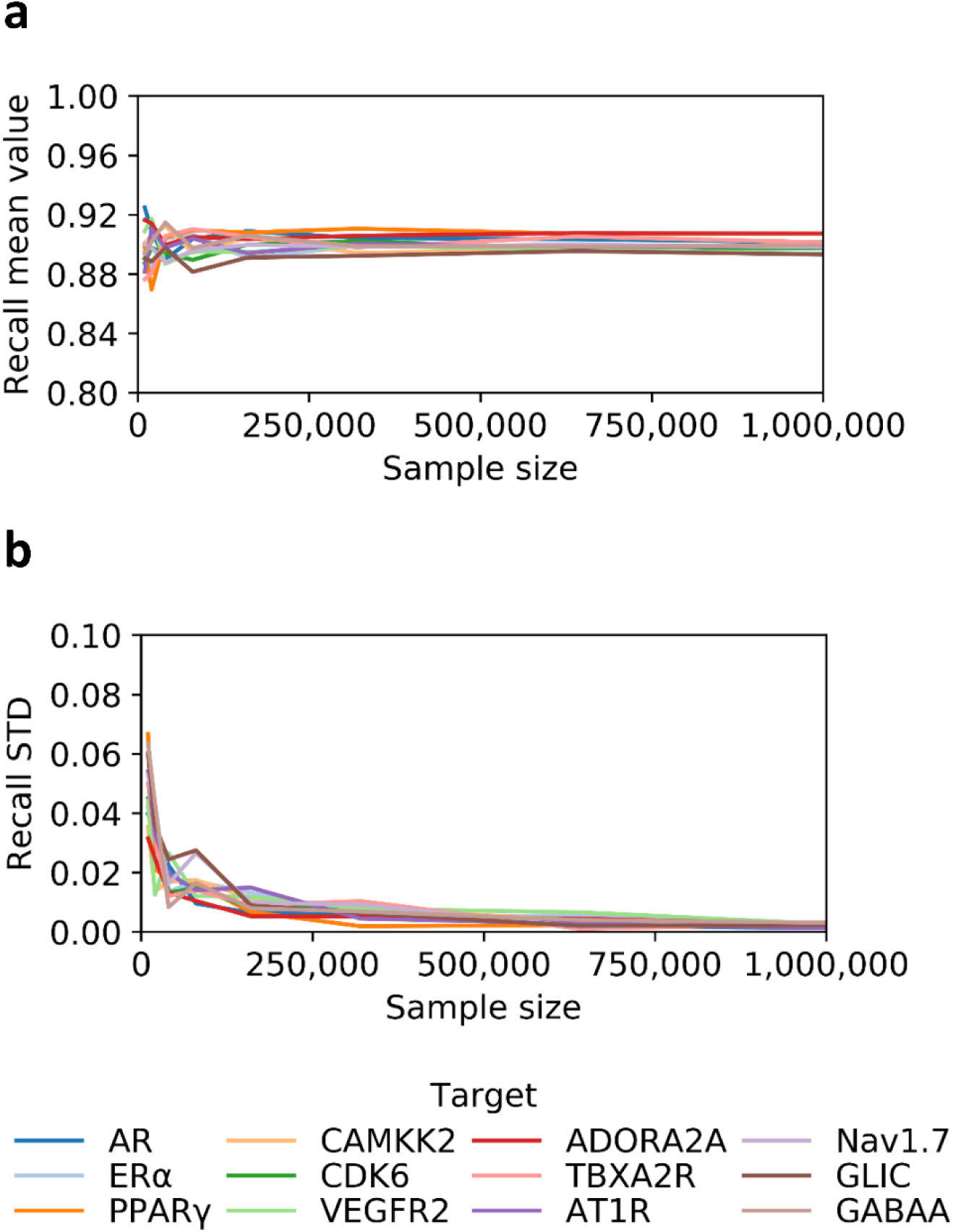
Effect of training set sample size on model generalizability. **a,** Mean values for the test set recalls computed using different sample sizes. Values approach 0.9 for all targets, when the training set size reaches about 1 million molecules. **b,** Similarly, variation of the standard deviations (STD) approach converts to 0, when the sample sizes reach 1 million molecules. We ran one iteration for each target and repeated computations 5 times for each sampling size. Androgen Receptor (AR), Estrogen Receptor-alpha (ERα), and Peroxisome Proliferator-Activated Receptor γ (PPARγ), Calcium/Calmodulin-Dependent Protein Kinase Kinase 2 (CAMKK2), Cyclin-Dependent Kinase 6 (CDK6), Vascular Endothelial Growth Factor Receptor 2 (VEGFR2), Adenosine A2A Receptor (ADORA2A), Thromboxane A2 Receptor (TBXA2R), Angiotensin II Receptor Type 1 (AT1R), Nav1.7 sodium channel (Nav1.7), Gloeobacter Ligand-Gated Ion Channel (GLIC), Gamma-aminobutyric acid receptor type A (GABAA).

### Size reduction of ZINC15 by *D*^*2*^ virtual screening

The main goal of *D*^*2*^ methodology is to reduce billions of Big Base entries to a manageable few-million-molecules subset which yet encompasses the vast majority of potential virtual hits. This final ranked molecular subset can then be normally docked into the target using one or several docking programs, or can post-processed with other VS means. The *D*^*2*^ method relies on iterative improvement of the deep neural network (DNN) training set by adding predicted hit molecules from each previous iteration, while the deciding cutoff also gradually becomes more stringent. We extensively evaluated the performance of this *D*^*2*^ protocol by screening all 1.36 billion molecules from ZINC15 against the 12 protein targets introduced above, using docking program FRED^16^.

For each target, we ran a total of 11 *D*^*2*^ iterations -one initial training step (requiring docking of 3 million entries) and 10 consecutive iterative docking steps, each involving docking of 1 million molecules. Thus, for each target we practically docked only 13 million molecular structures representing only ~1% of the 1.36 billion entries of ZINC15. Figure 3a illustrates the docking score cutoffs used to discern hits and non-hits at each iteration. These values decreased in accordance with the lowering of percentage of molecules defined as virtual hits in the validation sets at each iteration (see ‘Methods’ section for details).

**Figure 3.**
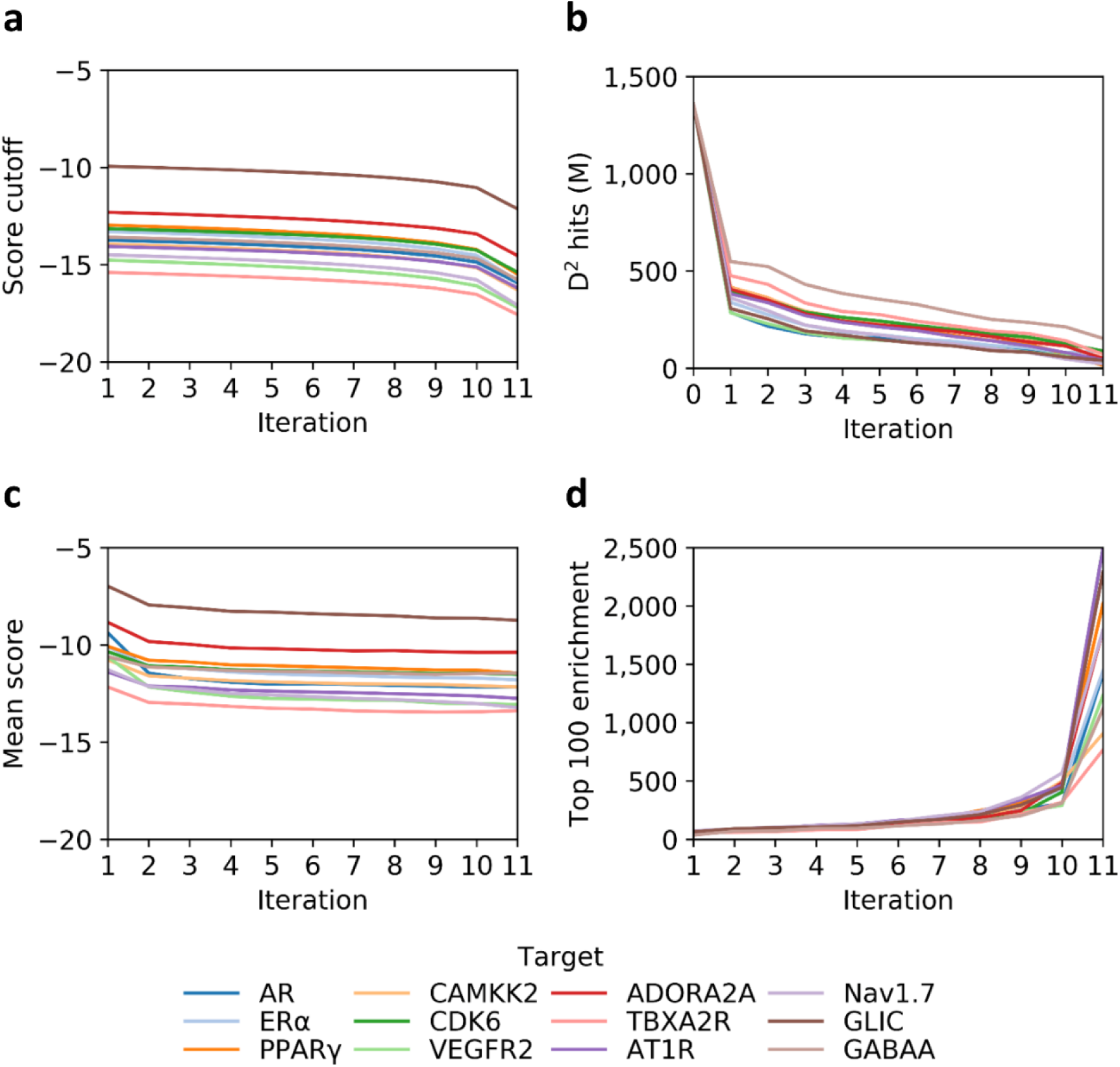
D^2^ performance statistics for 12 drug targets. **a,** Variation of score cutoff values used for selecting virtual hits at each iteration. **b,** Variation of numbers of molecules predicted as virtual hits after each iteration. **c,** Iterative improvement of docking score mean values for molecules randomly selected for training set augmentation. **d,** Enrichment values calculated for 100 top ranked predicted virtual hits in test set after each iteration. Androgen Receptor (AR), Estrogen Receptor-alpha (ERα), Peroxisome Proliferator-Activated Receptor γ (PPARγ), Calcium/Calmodulin-Dependent Protein Kinase Kinase 2 (CAMKK2), Cyclin-Dependent Kinase 6 (CDK6), Vascular Endothelial Growth Factor Receptor 2 (VEGFR2), Adenosine A2A Receptor (ADORA2A), Thromboxane A2 Receptor (TBXA2R), Angiotensin II Receptor Type 1 (AT1R), Nav1.7 sodium channel (Nav1.7), Gloeobacter Ligand-Gated Ion Channel (GLIC), Gamma-aminobutyric acid receptor type A (GABAA).

The majority of non-hits was removed during the first iteration for all targets, while less molecules were discarded in successive steps, as expected due to larger portions of unfavorable compounds being present at the beginning of the runs. The decrease rate and the number of hits identified were target-dependent (Figure 3b).

Another notable observation from the analysis of the *D*^*2*^ progression is that training sets are effectively improved after each iteration, as indicated by the mean docking scores of samples added to training that shifted toward more negative (favorable) values after each round of modeling (Figure 3c). Importantly, this observation marks progressively more confident performance of *D*^*2*^ in recognizing and discarding unfavorable molecular structures, and consequently favorably augmenting the training set. Thus, we anticipated *D*^2^ to improve the enrichment of predicted subsets for virtual hits after each iteration, as consequence of augmenting procedure. Figure 3d features the resulting trends of enrichment values for the top 100 molecules in the test sets ranked by the DNN models. As data indicate, these values increased after each iteration for all targets, also suggesting that model’s performance improves every time the training set is augmented with molecules from each previous *D*^*2*^ iterative step. Of note, the enrichment values strongly increased in the last iteration, where the models retrieved a very small portion (0.01%) of the top scoring molecules of the Big Base. Further indications of iterative improvement of DNN models are provided by the area under the curve receiver operating characteristics (AUC ROC) values and full predicted database enrichment (FPDE) values, presented in Supplementary Table S2 for all 12 studied targets.

### Analysis of *D*^*2*^ performance

As indicated earlier the main objective of the *D*^*2*^ methodology is to reduce a Big Base to a highly enriched library of molecules that can be processed using conventional docking programs and computational resources in order to remove the remaining unfavorable entries. We observed that the sizes of the final subset ranged between 1% and 12% of the original Big Base, depending on the target (Figure 4a). It is foreseen that these remaining enriched and DNN-ranked libraries can then be post-processed to remove residual low scoring molecules. Alternatively, *D*^*2*^ can be carried out until the convergence of a Big Base.

**Figure 4.**
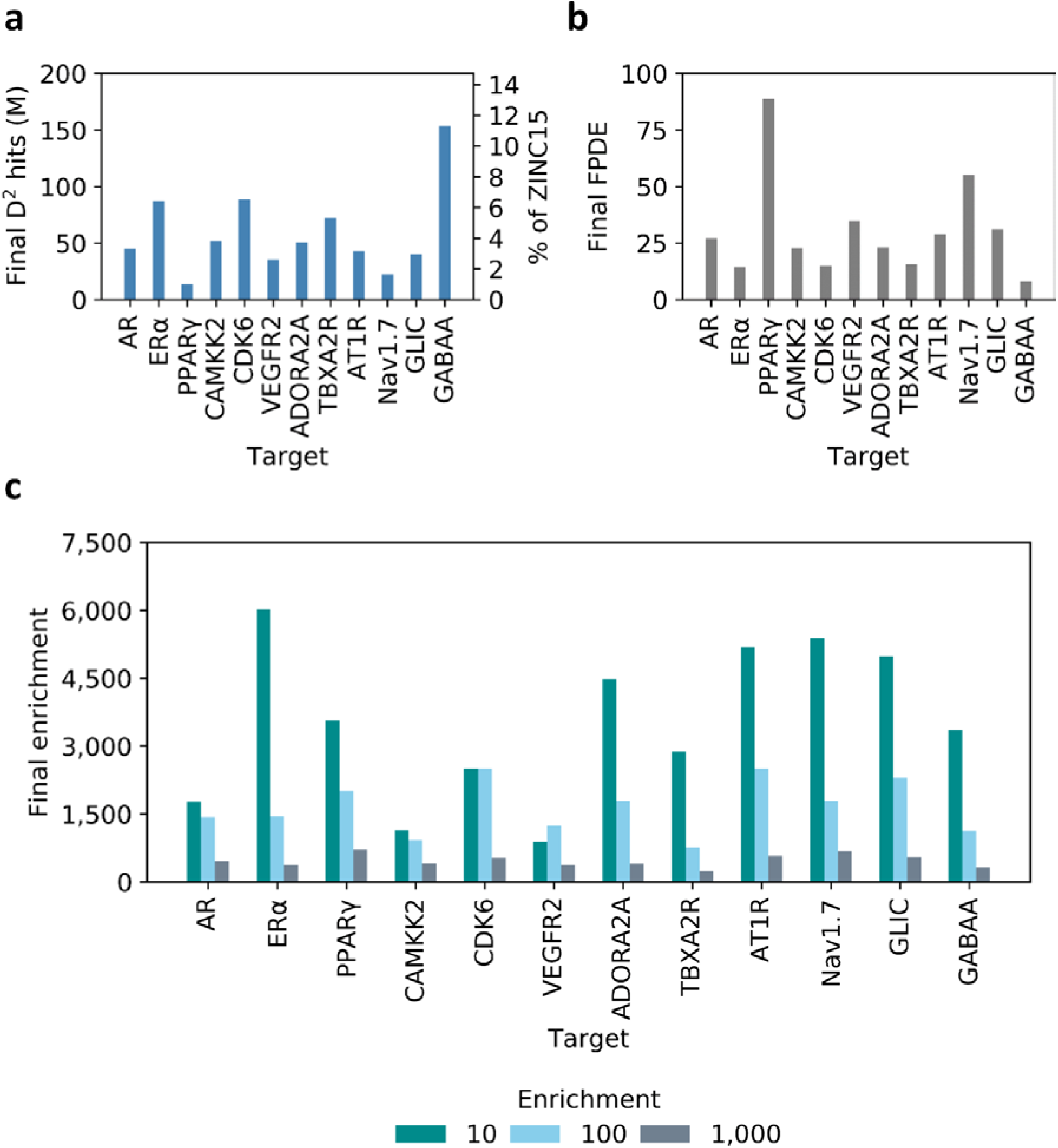
Final dataset sizes and enrichment values resulting from D^2^ runs. **a,** Total number of molecules predicted as virtual hits remaining after the eleventh D^2^ iteration. Values are also reported in terms of percentage of ZINC15 entries that were retained (right vertical axis). **b,** Full predicted database enrichment (FPDE) values resulting from the last iterations of D^2^ experiments. **c,** Enrichment values for top 10, top 100 and top 1,000 selected virtual hits (in the test sets). Androgen Receptor (AR), Estrogen Receptor-alpha (ERα), Peroxisome Proliferator-Activated Receptor γ (PPARγ), Calcium/Calmodulin-Dependent Protein Kinase Kinase 2 (CAMKK2), Cyclin-Dependent Kinase 6 (CDK6), Vascular Endothelial Growth Factor Receptor 2 (VEGFR2), Adenosine A2A Receptor (ADORA2A), Thromboxane A2 Receptor (TBXA2R), Angiotensin II Receptor Type 1 (AT1R), Nav1.7 sodium channel (Nav1.7), Gloeobacter Ligand-Gated Ion Channel (GLIC), Gamma-aminobutyric acid receptor type A (GABAA).

Overall, *D*^*2*^ demonstrated its best performance for PPARγ protein, for which the size of ZINC15 was reduced to 1%. Thus, for this system, the total number of docked molecules for screening ZINC15 was 50 times less than conventional VS, including molecules required for model training and molecules remaining after the eleventh iteration. On the other hand, *D*^*2*^ was the least effective on the GABAA system, where 12% of ZINC15 molecules were left after the last iteration. Encouragingly, the recall values were consistently transferred to the test sets for all *D*^*2*^ iterations for all targets, indicating that all underlying DL models were generalizable in a consistent way (see Supplementary Table S2). To further assess the overall *D*^*2*^ performance we evaluated FPDE and enrichment values for the top 10, 100 and 1,000 virtual hits generated in the test sets after the final iteration, resulting in enrichment values ranging from 240 to 6,000 as demonstrated on Figure 4b, and FPDE values ranging from 8 to 88 (Figure 4b).

Is important to note, that we have set stringent 90% recall values for *D*^*2*^ iterations to preserve the vast majority of potential virtual hits in a final molecular set. However, one can relax such recall cutoff to lower values to sacrifice the retention of virtual hits in a *D*^*2*^ workflow, but to significantly reduce the number of actually docked molecules and to shorten the *D*^*2*^ runtime.

Overall, the above analysis clearly indicates that *D*^*2*^ procedure can effectively discard most of unqualified molecules in a Big Base, without losing more than a predefined percentage of virtual hits (10% in the case of the current study). In our opinion, this makes *D*^*2*^ methodology an efficient mean for conducting large-scale VS campaigns involving billions of small molecule structures, and a valid alternative to brute force approaches demanding large amounts of computational resources.

## CONCLUSIONS

With the increasing automation of synthetic procedures, the focus of modern drug discovery campaigns will be shifting towards screening of increasingly larger molecular libraries consisting of billions of chemical structures. To reinforce such opportunity, docking protocols demonstrate improved performance on large make-on-demand databases, effectively yielding novel, diverse and nontraditional chemotypes for drug discovery endeavors^6^. It could be noted, however, that most of time and resources invested in modern docking campaigns are spent on processing unfavorable molecular structures, while the emerging ‘negative’ data is also not being utilized in any way or form.

Hence, to keep the pace with ever-expanding chemical Big Data space and to fully utilize results generated by docking programs ‘on a fly’, we have developed the *Deep Docking* protocol *D*^*2*^, a DNN-based method for processing very large chemical libraries with conventional resources. The method relies on iterative docking of a small portion of a parental Big Base (such as ZINC15) and utilizes generated docking scores (both favorable and unfavorable) to train ligand-based QSAR models. These models then enable to approximate docking outcome for unprocessed Big Base entries. We have demonstrated that such approach can yield a manageable small subset of a Big Base, highly enriched with favorably ‘dockable’ molecular structures.

We demonstrated the power of *D*^*2*^ by screening all 1.36 billion entries of ZINC15 against 12 prominent drug targets, where the original Big Base was significantly reduced while retrieving a controlled, high portion of favorably docked molecules. At the same time, most of low scoring molecules were removed without investing time and resources in them and the generated ZINC subsets were highly enriched in potential virtual hits. Notably, screening a Big Base using *D*^*2*^ requires to dock up to 50 times fewer molecules compared to conventional docking, while losing only about 10% of virtual hits.

Collectively, the results reported in this study strongly advocate the use of *D*^*2*^ for large-scale screening campaigns, as it eliminates the need of processing *a priori* unfavorable molecular structures and provides significant savings of processing time required for docking billions of molecules.

## METHODS

### QSAR descriptors

SMILES of 1.36 billion molecular structures were downloaded from ZINC15^5^. Morgan fingerprints with a size of 1024 bits and a radius of 2 were generated using the RDKit package^17^.

### Protein targets

The x-ray structures of AR^18^, ERα^19^, PPARγ^20^, CAMKK2^21^, CDK6^22^, VEGFR2^23^, ADORA2A^24^, TBXA2R^25^, AT1R^26^, Nav1.7^27^, GLIC^28^, and GABAA^29^ containing co-crystallized ligands were extracted from the Protein Data Bank (PDB)^30^. Details about the selected target structures are summarized in Supplementary Table S2.

### Molecular Docking

PDB structures were optimized using the Protein Preparation Wizard module from Schrödinger suite^31^, and docking grids were prepared using the MakeReceptor utility from OpenEye^32^. SMILES were processed using QUACPAC^33^ in order to generate dominant tautomer and ionization states at pH 7.4. The OMEGA’s pose module^34, 35^ was used to generate 3D conformers for docking. Docking simulations were carried out using FRED program and Chemgauss4 scoring function from OpenEye^16^.

### Database sampling

The optimal number of molecules required for the training set was determined by running one *D*^*2*^ iteration for each target, using different sizes for training, validation and test set (10,000, 20,000, 40,000, 80,000, 160,000, 320,000, 640,000 and 1 million molecules). For each sample size, computations were repeated 5 independent times. The optimal sampling size was then chosen by evaluating means and standard deviations of recall values in the test sets for all targets.

### *D*^*2*^ workflow

Initial training, validation and test sets used for the DL model consisted of 1 million molecules each that were randomly sampled from ZINC15 during the first *D*^*2*^ iteration. Each set was docked to the target of interest and then divided into virtual hits and non-hits based on the generated docking scores. The score cutoff used to determine the class of molecules was determined in order to split the validation in 1% top scoring molecules (virtual hits) and 99% non-hits. Molecules of each set with docking scores equally or more favorable than the cutoff value were assigned to the virtual hit class, while remaining molecules were assigned to the non-hit class. The DNN model was trained using classes and molecular descriptors of the processed entries, and then used to predict virtual hits and unqualified molecules from the whole ZINC15 based on molecular descriptors. From the second iteration onward, the training set was expanded with 1 million molecules randomly sampled from hits predicted in the previous iteration. The score cutoff was gradually decreased (corresponding to higher predicted target affinity) after each iteration to keep selecting better compounds. This reduction was done by linearly lowering the percentage of top scoring molecules in the validation set assigned to the virtual hit class from 1% in the first iteration to 0.01% in the last one. Thus, the cutoff value in iteration 2 corresponded to the highest (worst) docking score of the top 0.9% ranked compounds, in iteration 3 it corresponded to the highest docking score of top 0.8% compounds, and so on.

### Evaluation metrics

All evaluation metrics were calculated on the test sets. Precision was calculated as:

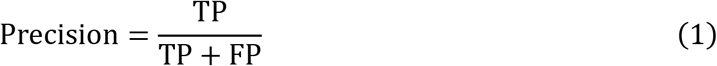

where TP (true positives) were virtual hits correctly predicted by the DNN, and FP (false positives) were actual non- hits that were incorrectly classified as virtual hits by the DNN.

Recall was calculated as:

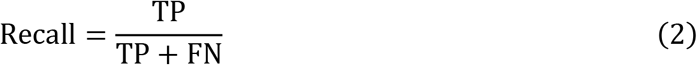

where FN (false negatives) were virtual hits incorrectly discarded by the DNN. Enrichment values were calculated as:

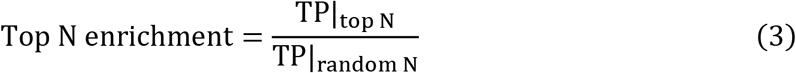

with TP|_top N_ as the number of TP found within the top N ranked molecules by the DNN, and TP|_random N_ as the number of TP found within N randomly sampled molecules. N was set to values equal to 10, 100 and 1,000.

FPDE was calculated as the ratio between precision (Equation 1) and random precision:

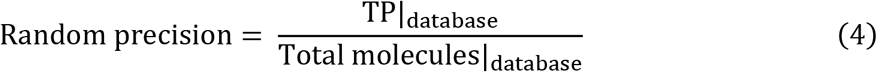

### Deep learning

The Keras Python library^36^ was used for building and training feed-forward DNN models^37^. Model hyperparameters were set as number of hidden layers and neurons, dropout frequency, as well as oversampling of the minority class and class weights, in order to deal with highly imbalanced datasets (1% to 0.01% of virtual hits). A lower threshold value was established for the DNN probabilities (indicating the likelihood of molecules of being virtual hits), and used as criterion to assign molecules to the virtual hit class upon prediction. The threshold was chosen each time in order to retrieve 90% of the actual virtual hits (i.e. top scoring molecules) of the validation set. Model selection was performed by running a basic grid search to identify the set of hyperparameters providing the highest FPDE value in the test set. The best model was then applied to all ZINC15 entries in order to predict virtual hits and non-hits.

### Hardware

We used 6 Intel(R) Xeon(R) Silver 4116 CPU @ 2.10GHz (a total of 60 cores) for docking, and 4 Nvidia Tesla V100 GPUs with 32GB memory for DNN model training and inference.

## CODE AVAILABILITY

We made the *D*^*2*^ code publicly available in GitHub at https://github.com/vibudh2209/D2.

## DATA AVAILABILITY

Docking datasets used for building models are available at https://drive.google.com/file/d/1w86NqUk7brjDIGCxD65tFLNeQ5IgLeHZ/view?usp=sharing.

## ACKNOWLEDGMENTS

This work has been supported by the Canadian Cancer Society Research Institute (CCSRI) Impact Grant 2019 #706145.

