## Supplementary material for "Deep Docking - a Deep Learning Approach for Virtual Screening of Big Chemical Datasets"

***Table S1. Protein targets selected for evaluating D$^2$.***

| **Family** | **Target** | **Ligand** | **PDB** | **Resolution (Å)** | **Therapeutic Significance** |
|---|---|---|---|---|---|
| Nuclear receptors | AR | Dihydrotestosterone | 1T7R[1] | 1.40 | Prostate Cancer[2] |
| | ERα | Raloxifene | 1ERR[3] | 2.60 | Breast cancer, osteoporosis, menopausal symptoms[2] |
| | PPARγ | SR-2067 | 4R06[4] | 2.22 | Diabetes[2] |
| Kinases | CAMKK2 | STO-609 | 2ZV2[5] | 2.40 | Prostate cancer, metabolic hepatic diseases[5, 6] |
| | CDK6 | Abemaciclib | 5L2S[7] | 2.27 | Breast cancer[8] |
| | VEGFR2 | Axitinib | 4AG8[9] | 1.95 | Multiple cancer types[8] |
| G protein-coupled receptors | ADORA2A | Theophylline | 5MZJ[10] | 2.00 | Myocardial perfusion imaging, inflammation, neuropathic pain, Parkinson’s disease[11] |
| | TBXA2R | Ramatroban | 6IIU[12] | 2.50 | Cardiovascular diseases, asthma[13] |
| | AT1R | ZD-7155 | 4YAY[14] | 2.90 | Hypertension[15] |
| Ion channels | Nav1.7 | GX-936 | 5EK0[16] | 3.53 | Pain[17] |
| | GLIC | Anesthetic ketamine | 4F8H[18] | 2.99 | General anesthetics[18] |
| | GABAA | Flumazenil | 6D6T[19] | 3.86 | Benzodiazepine overdose[19] |

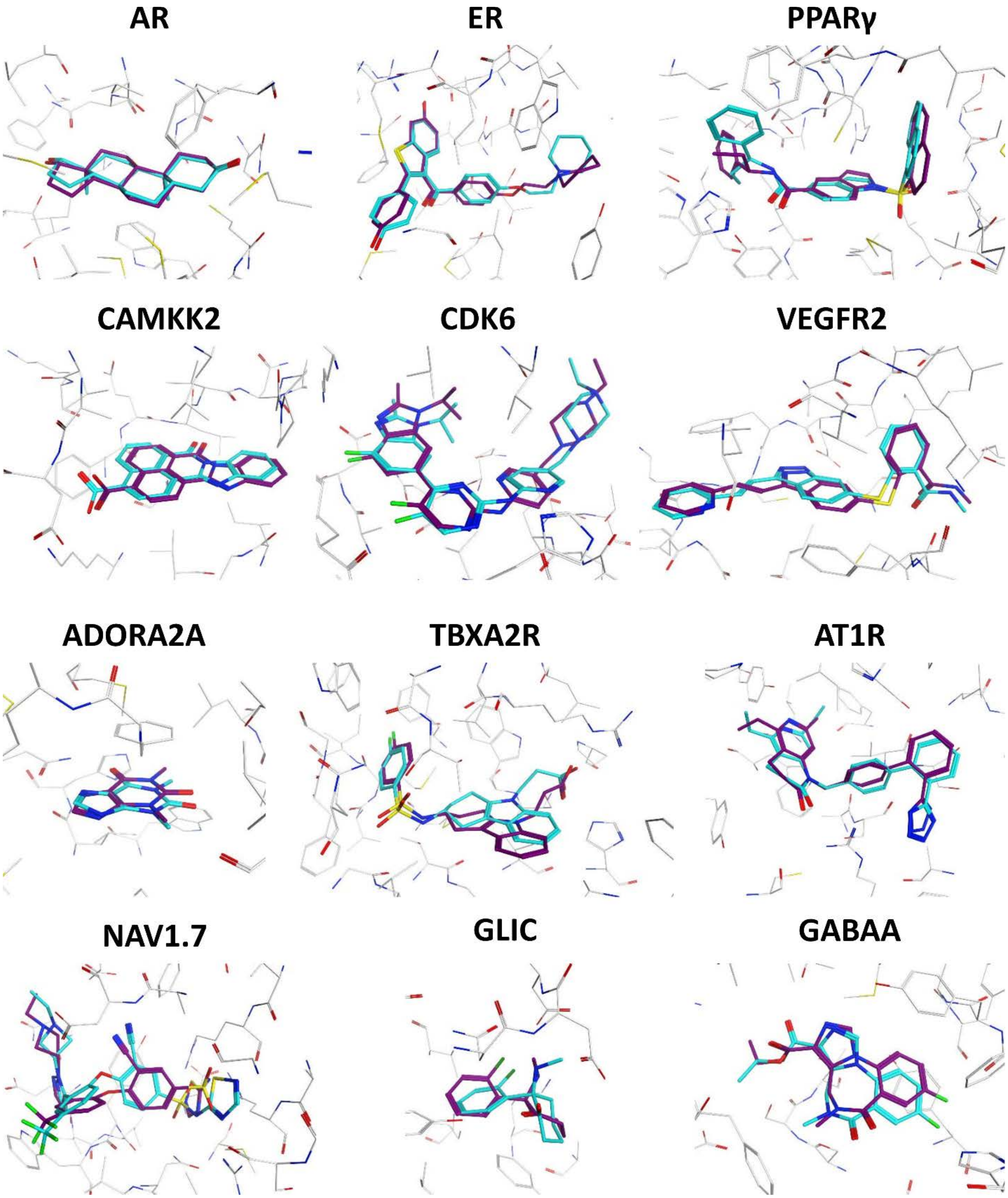


***Figure S1. Results of FRED docking of native ligands to their receptor structures.***

*Superposition of x-ray binding poses (cyan carbons) and docking poses (purple carbons) of the ligands for the 12 investigated systems.*

*Table S3. Performance of $D^2$ at each iteration.*

| Target | Iteration | FPDE | ROC AUC | Test set recall (%) |
|---|---|---|---|---|
| AR | 1 | 4.31 | 0.93 | 0.90 |
| | 2 | 5.68 | 0.94 | 0.89 |
| | 3 | 6.92 | 0.95 | 0.89 |
| | 4 | 7.80 | 0.96 | 0.89 |
| | 5 | 7.93 | 0.96 | 0.90 |
| | 6 | 8.83 | 0.96 | 0.90 |
| | 7 | 9.67 | 0.97 | 0.90 |
| | 8 | 10.81 | 0.97 | 0.89 |
| | 9 | 12.69 | 0.97 | 0.89 |
| | 10 | 15.92 | 0.97 | 0.89 |
| | 11 | 27.17 | 0.98 | 0.88 |
| ERα | 1 | 3.62 | 0.91 | 0.90 |
| | 2 | 4.44 | 0.93 | 0.90 |
| | 3 | 5.54 | 0.94 | 0.90 |
| | 4 | 6.49 | 0.95 | 0.89 |
| | 5 | 7.06 | 0.95 | 0.89 |
| | 6 | 8.02 | 0.96 | 0.89 |
| | 7 | 8.98 | 0.96 | 0.91 |
| | 8 | 10.29 | 0.97 | 0.89 |
| | 9 | 11.83 | 0.97 | 0.90 |
| | 10 | 15.32 | 0.98 | 0.90 |
| | 11 | 14.36 | 0.97 | 0.93 |
| PPARγ | 1 | 2.95 | 0.90 | 0.90 |
| | 2 | 3.39 | 0.92 | 0.91 |
| | 3 | 4.31 | 0.93 | 0.90 |
| | 4 | 4.81 | 0.94 | 0.90 |
| | 5 | 5.42 | 0.95 | 0.90 |
| | 6 | 6.23 | 0.95 | 0.90 |
| | 7 | 7.22 | 0.96 | 0.90 |
| | 8 | 8.54 | 0.96 | 0.90 |
| | 9 | 9.90 | 0.97 | 0.90 |
| | 10 | 16.44 | 0.98 | 0.89 |
| | 11 | 88.71 | 1.00 | 0.93 |
| CAMKK2 | 1 | 2.95 | 0.89 | 0.90 |
| | 2 | 3.40 | 0.91 | 0.90 |
| | 3 | 4.14 | 0.93 | 0.90 |
| | 4 | 4.70 | 0.94 | 0.90 |
| | 5 | 5.15 | 0.94 | 0.90 |
| | 6 | 5.64 | 0.95 | 0.90 |
| | 7 | 6.13 | 0.95 | 0.91 |
| | 8 | 6.78 | 0.96 | 0.90 |
| | 9 | 7.95 | 0.96 | 0.90 |

| | | | | |
|---|---|---|---|---|
| | 10 | 10.77 | 0.97 | 0.91 |
| | 11 | 22.93 | 0.98 | 0.87 |
| CDK6 | 1 | 3.13 | 0.90 | 0.90 |
| | 2 | 3.51 | 0.91 | 0.90 |
| | 3 | 4.25 | 0.93 | 0.90 |
| | 4 | 4.68 | 0.94 | 0.90 |
| | 5 | 5.03 | 0.94 | 0.90 |
| | 6 | 5.55 | 0.94 | 0.90 |
| | 7 | 6.18 | 0.95 | 0.90 |
| | 8 | 6.90 | 0.95 | 0.90 |
| | 9 | 7.66 | 0.96 | 0.90 |
| | 10 | 9.79 | 0.97 | 0.90 |
| | 11 | 15.10 | 0.99 | 0.99 |
| VEGFR2 | 1 | 4.28 | 0.93 | 0.90 |
| | 2 | 5.33 | 0.94 | 0.90 |
| | 3 | 6.66 | 0.95 | 0.90 |
| | 4 | 7.79 | 0.96 | 0.89 |
| | 5 | 8.41 | 0.96 | 0.90 |
| | 6 | 9.21 | 0.97 | 0.90 |
| | 7 | 10.17 | 0.97 | 0.90 |
| | 8 | 11.98 | 0.97 | 0.91 |
| | 9 | 14.68 | 0.98 | 0.90 |
| | 10 | 18.76 | 0.98 | 0.90 |
| | 11 | 34.85 | 0.97 | 0.91 |
| ADORA2A | 1 | 3.01 | 0.90 | 0.90 |
| | 2 | 3.52 | 0.92 | 0.90 |
| | 3 | 4.38 | 0.93 | 0.90 |
| | 4 | 5.02 | 0.94 | 0.89 |
| | 5 | 5.48 | 0.94 | 0.89 |
| | 6 | 5.93 | 0.95 | 0.90 |
| | 7 | 6.54 | 0.95 | 0.90 |
| | 8 | 7.56 | 0.95 | 0.90 |
| | 9 | 9.03 | 0.96 | 0.90 |
| | 10 | 10.79 | 0.97 | 0.90 |
| | 11 | 23.05 | 0.97 | 0.85 |
| TBXA2R | 1 | 2.58 | 0.88 | 0.90 |
| | 2 | 2.84 | 0.89 | 0.90 |
| | 3 | 3.62 | 0.91 | 0.89 |
| | 4 | 4.16 | 0.92 | 0.89 |
| | 5 | 4.45 | 0.93 | 0.90 |
| | 6 | 5.07 | 0.94 | 0.90 |
| | 7 | 5.64 | 0.94 | 0.90 |
| | 8 | 6.41 | 0.95 | 0.90 |
| | 9 | 6.91 | 0.95 | 0.91 |
| | 10 | 8.79 | 0.96 | 0.91 |

| | | | | |
|---|---|---|---|---|
| | 11 | 15.61 | 0.96 | 0.83 |
| AT1R | 1 | 3.20 | 0.90 | 0.90 |
| | 2 | 3.63 | 0.92 | 0.90 |
| | 3 | 4.51 | 0.93 | 0.90 |
| | 4 | 5.16 | 0.94 | 0.90 |
| | 5 | 5.75 | 0.95 | 0.90 |
| | 6 | 6.30 | 0.95 | 0.90 |
| | 7 | 7.49 | 0.96 | 0.90 |
| | 8 | 8.63 | 0.96 | 0.90 |
| | 9 | 10.63 | 0.97 | 0.90 |
| | 10 | 15.27 | 0.98 | 0.91 |
| | 11 | 28.95 | 0.98 | 0.91 |
| Nav1.7 | 1 | 3.39 | 0.91 | 0.90 |
| | 2 | 4.15 | 0.93 | 0.90 |
| | 3 | 5.50 | 0.94 | 0.90 |
| | 4 | 6.37 | 0.95 | 0.90 |
| | 5 | 7.15 | 0.96 | 0.90 |
| | 6 | 8.37 | 0.96 | 0.89 |
| | 7 | 9.69 | 0.96 | 0.89 |
| | 8 | 11.99 | 0.97 | 0.89 |
| | 9 | 15.02 | 0.98 | 0.89 |
| | 10 | 25.53 | 0.98 | 0.89 |
| | 11 | 55.26 | 0.98 | 0.90 |
| GLIC | 1 | 3.95 | 0.92 | 0.89 |
| | 2 | 4.82 | 0.94 | 0.91 |
| | 3 | 6.35 | 0.95 | 0.90 |
| | 4 | 7.13 | 0.96 | 0.90 |
| | 5 | 8.31 | 0.96 | 0.90 |
| | 6 | 9.45 | 0.97 | 0.90 |
| | 7 | 10.61 | 0.97 | 0.90 |
| | 8 | 13.35 | 0.97 | 0.89 |
| | 9 | 14.84 | 0.98 | 0.90 |
| | 10 | 21.32 | 0.98 | 0.90 |
| | 11 | 31.03 | 0.98 | 0.92 |
| GABAA | 1 | 2.23 | 0.86 | 0.90 |
| | 2 | 2.34 | 0.87 | 0.90 |
| | 3 | 2.83 | 0.89 | 0.90 |
| | 4 | 3.17 | 0.91 | 0.90 |
| | 5 | 3.45 | 0.91 | 0.90 |
| | 6 | 3.74 | 0.92 | 0.90 |
| | 7 | 4.25 | 0.93 | 0.90 |
| | 8 | 4.80 | 0.93 | 0.89 |
| | 9 | 5.13 | 0.94 | 0.89 |
| | 10 | 5.80 | 0.95 | 0.90 |
| | 11 | 8.05 | 0.94 | 0.91 |
